# Creative destruction: Sparse activity emerges on the mammal connectome under a simulated communication strategy with collisions and redundancy

**DOI:** 10.1101/2020.02.25.964593

**Authors:** Yan Hao, Daniel Graham

**Affiliations:** Department of Mathematics and Computer Science, Hobart & William Smith Colleges Geneva, NY 14456 USA; Department of Psychology, Hobart & William Smith Colleges Geneva, NY 14456 USA

**Keywords:** connectome, communication systems, collisions, redundancy, routing, sparseness

## Abstract

Signal interactions in brain network communication have been little studied. We describe how nonlinear collision rules on simulated mammal brain networks can result in sparse activity dynamics characteristic of mammalian neural systems. We tested the effects of collisions in “information spreading” (IS) routing models and in standard random walk (RW) routing models. Simulations employed synchronous agents on tracer-based mesoscale mammal connectomes at a range of signal loads. We find that RW models have high average activity that increases with load. Activity in RW models is also densely distributed over nodes: a substantial fraction is highly active in a given time window, and this fraction increases with load. Surprisingly, while IS models make many more attempts to pass signals, they show lower net activity due to collisions compared to RW, and activity in IS increases little as function of load. Activity in IS also shows greater sparseness than RW, and sparseness decreases slowly with load. Results hold on two networks of the monkey cortex and one of the mouse whole-brain. We also find evidence that activity is lower and more sparse for empirical networks compared to degree-matched randomized networks under IS, suggesting that brain network topology supports IS-like routing strategies.

## Introduction

How does the mammal brain manage the communication of signals across its network? At the level of mesoscale mammal brain network structure, comprising connectivity of tens to hundreds of brain areas over thousands to hundreds of thousands of connections, an increasingly detailed picture of network connectivity is emerging (Markov et al., 2014; Oh et al., 2016; Gamanut et al., 2018). Researchers are now aiming to “animate” these networks in order to understand patterns of whole-brain signaling dynamics (a research area termed *dynomics*; Kopell et al., 2014).

However, a given topology can support a wide range of observable dynamics (see, e.g., Friston, 2011; Knoblauch et al., 2016). Adding to the challenge is that activity on the whole network is poorly understood, with severe limitations in sampling at the single neuron level, and in spatial and temporal precision for imaging methods. Moreover, though single-neuron spiking dynamics for some neuron classes are well understood, global activity will not necessarily be a linear summation of single-neuron dynamics.

One approach to understanding brain network dynamics is to consider the system of intercommunication among nodes as a bridge between network topology and network dynamics. Communication dynamics can serve as a linkage between structural connectivity and functional connectivity (Avena-Koenigsberger et al., 2018). More generally, nodes in the network can be considered as communicators that attempt to pass signals or messages to other nodes according to routing rules (Graham, 2014).

Communication demands such as flexibility and reliability place important constraints on the system (Poggio, 1984; Hahn et al., 2018). Effective and robust routing strategies are especially important on the mammal connectome because the topology is such that paths of just a few synapses exist between any node and practically any other node. Brain networks also need to dynamically exchange multimodal information among a large number of nodes in real time without appreciably changing network topology. In addition, routing in the brain lacks central control.

To begin to understand what routing strategy is in use in the brain, one can look to design strategies in engineered systems (Graham and Rockmore, 2010; Graham 2014; 2017; Navlakha et al., 2015; 2017; Fornito et al., 2016). A fundamental engineering goal for any large-scale communication system is the management of signal interactions. Signal interactions could take many forms such as summation, thresholding, collision, and duplication. As we describe in the Discussion, brain-like networks with signal interactions involving summations/thresholds tend to produce undesirable behavior such as activity die-off or overload (Kaiser et al., 2007; Kaiser and Hilgetag, 2010). Here we perform initial simulations of two interactions that have received little attention: signal collision and signal duplication (i.e., redundancy).

Collisions occur when signals from different sources converge on the same target at the same time. They are the price paid for the ability to route signals selectively to many possible destinations on a fixed network. Collision dynamics are emergent: they are a nonlinear effect of topology, node dynamics, and current traffic. All large-scale engineered communication systems have means for managing collisions, through redress, arbitration, and other strategies (see e.g., Kleinrock, 1976; Mišic and Mišic, 2014).

Given the *a priori* likelihood of collisions on small world-like brain networks (compared to, for example, lattice networks), we argue that it is in the interest of brain networks to establish dynamic interactions that manage collisions successfully across the entire brain. Management of collisions is especially important in mesoscale brain networks since behavior-related information must be exchanged among a large number of subsystems.

However, collisions introduce nonlinearities, which are difficult to study analytically. Nonlinear signal interactions are increasingly investigated with explicit agent-based models in the study of other communication networks (e.g., epidemics: Min et al. 2020), but little work in network neuroscience has used explicit approaches. For example, most random walk models (e.g., Noh and Rieger, 2004; Abdelnour et al., 2014), which route signals to randomly chosen outgoing edges use analytical methods that cannot account for signal interactions such as collisions.

An exception is Mišić et al. (2014a,b). In this pair of studies, buffers (node memory allocations, a form of nonlinearity) were employed to manage collisions in a random walk routing model. In this scheme, colliding signals at the inputs of a node are lined up in a queue and stored in node memory until they can be directed to a randomly chosen outgoing edge. Buffers are ubiquitous on the Internet and may be useful in brains. But though several neurobiological mechanisms for buffering have been proposed, they remain hypothetical (see, e.g., Goldman-Rakic, 1996; Graham, 2014; Funahashi, 2015).

An alternative to random walk models is shortest path routing, which invokes the logic of evolved optimality (e.g., Bullmore and Sporns, 2009; Rubinov and Sporns, 2010; Goñi et al., 2013, 2014). Shortest path strategies also have parallels in engineered systems like the Internet (e.g., OSPF: open shortest-path first routing, see, e.g., Sosnovich et al., 2017). However, shortest path models typically ignore signal interactions such as collisions, again because these models are studied analytically. Yet signal interactions are especially important in a shortest path context. Though it has not been well recognized, successful shortest path routing requires knowledge of current network traffic: a short path is not necessarily short if there is congestion. Even when short path models include small buffers, they show substantial message loss due to collisions (Graham and Hao, 2018). In any case, shortest path routing is considered implausible because it requires global knowledge of network architecture to select shortest paths for all signals (Seguin et al., 2018; but see Mišić et al., 2015).

In the present work, we investigate a strategy of fully destructive collisions. The brain may in fact use a less punitive strategy than this but neurophysiological evidence increasingly supports the notion that signals in the brain regularly collide and are destroyed. For example, Sardi et al. (2017) demonstrated the failure of coincident signals above threshold to elicit spikes. They also showed that excitatory signals can cancel each other when they collide “head on” (Sardi et al., 2017; this is in fact predicted by the Hodgkin-Huxley model: Scott, 1977). Gidon et al. (2020) have shown exclusive-OR activity in single *ex vivo* human pyramidal cortical neurons, a behavior that effectively destroys some colliding signals. At the circuit level, gating circuits (e.g., Steriade and Paré, 2007; Gollisch and Meister, 2010), when closed, can also be conceived as destructive collisions between signals. If collisions need to be managed at the single cell level and the level of circuits, the same logic applies at the level of mesoscale brain networks (see Discussion).

Collisions need not incapacitate a communication network. They may be useful in establishing stable and robust dynamics with locally-implemented routing rules. In particular, we suggest that collisions could—almost paradoxically—help promote system efficiency.

First, destructive collisions may on average help ensure a low level of energy use. Energy efficiency is imperative given that the dynamic connectome of mammals has strong metabolic constraints (Bullmore and Sporns, 2012, Goñi et al., 2013). Destructive collisions may also promote homeostasis, which appears desirable for maintaining relatively constant activity in the face of changing system demands (e.g., Aeschbach et al., 1997; Turrigiano, 1999).

But the brain cannot achieve efficiency simply by using as little energy as possible (Poldrack, 2015). The brain instead operates in sparse fashion, in part because the sensory environment is sparse. To avoid confusion, we note that *sparseness* (or, equivalently, sparsity) in the present context refers to the distribution of activity across nodes and over time, rather than to the topology of the network (on a scale from “sparse” to “dense” connectivity). Sparse operation of a system is defined by the achievement of low average activity ratios (see, e.g., Földiák, 2002).

Although sparse activity can confer low average energy usage, it does not necessarily imply minimal energy use. Instead, it involves a characteristically non-Gaussian distribution of activity across units, in particular a distribution with a strong peak near zero activity (most units are “off” at a given time) and heavy tails (a small number of units are likely to be highly active).

In the brain, it has been understood for decades that only a small fraction of units—10% or less—in a given part of the network can be highly active at a given time (Levy and Baxter 1997; Lennie, 2000; Attwell and Laughlin, 2003); this is known as population sparseness. In addition, it is estimated that a unit can only be highly active over a small fraction of its lifetime; this is known as lifetime sparseness (see Willmore and Tolhurst, 2001; Graham and Field, 2006). Sparse activity in the visual system, for example, is thought to be in part a result of sparse physical environments (Field, 1994; Olshausen and Field 1996; Bell and Sejnowski, 1997), in addition to metabolic constraints and other factors (Graham and Field, 2009). We suggest here that collision dynamics that result in emergent sparse activity are good candidates for how the mammal brain routes information.

If collisions are destructive, it may be necessary for the brain to adopt strategies to generate redundancy in order to promote reliability. Brain network communication models increasingly include not just a pathfinding dimension (spanning a spectrum from random walks to shortest paths: Avena-Koenigsberger et al., 2019), and also a second dimension that can be seen as redundancy (Avena-Koenigsberger et al., 2017; Bettinardi et al., 2017; Tipnis et al., 2018).

A simple way to promote redundancy is to produce redundant copies of messages at each node. Based on this idea, we introduce an *information spreading* (IS) model of brain network communication. Signal interactions in a system with collisions and information spreading will produce nonlinear, emergent behavior. For example, as more signals and copies of those signals propagate on the network, the influence of collisions is increasingly important. Using numerical simulations, we investigate emergent behavior of the IS model, in comparison to a standard random walk (RW) model. As described more fully in the Discussion section, IS and RW models should be seen as spanning a spectrum of distributed routing strategies, rather than as a test model and a control.

We show that mammal brain networks using IS achieve a balance between collisions and redundancy and they produce sparse activity. In particular, we show that, across a range of signal loads, low activity and high sparseness are emergent properties of network-wide communication under an IS model. Performance of the IS model contrasts with RW models, which exhibits high and increasing energy use with increasing signal loads, and lower and decreasing sparseness across loads. To investigate whether empirical topology in the mammal connectome supports sparse, efficient activity patterns, we also compare dynamics of the anatomical network to randomized networks with matched degree distributions.

## Methods

### Overview

Using a Markovian agent-based model on monkey and mouse tracer-based connectomes, we implemented an information spreading (IS) model wherein nodes pass copies of incoming signals to all outgoing edges. Simulations utilized a nonlinear collision rule whereby all colliding messages are destroyed. We also tested a standard random walk (RW) model following the same collision rule. In RW, each message is passed from one node to another through a randomly chosen outgoing edge. At each time step, a set number of messages, which we term the *load* is injected to randomly chosen nodes; this is the primary independent variable in this study.

### Connectomes

We utilized three mesoscale mammal connectomes: two of the macaque monkey (Markov et al., 2014, termed *monkey1* in the present study as shorthand; and the CoCoMac database, see Bakker et al. 2012, termed *monkey2*), and one for the mouse (Oh et al., 2014, termed *mouse*). All edges are directed and have a weight of 1.

The *monkey1* connectome comprises 91 cortical nodes and 1615 edges, primarily in the visual system (ipsilateral). Of these nodes, 29 have in- and out-degree > 0, while the remainder are source nodes (in-degree = 0). This means that 62 nodes in *monkey1* are not reachable from the rest of the network, and only pass messages injected at those nodes. This limitation is a result of the retrograde tracing method used by Markov et al. (2014), and is the reason we included the *monkey2* network as well. For *monkey2*, we used only nodes that had an in-degree and an out-degree of at least 1. This set comprised nodes corresponding to 163 cortical regions and 5093 edges. The *mouse* connectome spans the entire mouse brain (ipsilateral), comprising 213 nodes and 16954 edges (all have in- and out-degree > 0). For all connectomes, isolated nodes with no incoming or outgoing edges (present only in *monkey2*) had been removed. We note that both *mouse* and *monkey2* contain self-loops (i.e., nodes with edges to themselves) but removing these edges did not change the outcome of the experiments so self-loops are included in the presented data. Including nodes with in-degree or out-degree of 0 also did not affect the outcome. Adjacency matrices for the data sets are shown in Figure 1.

**Figure 1.**
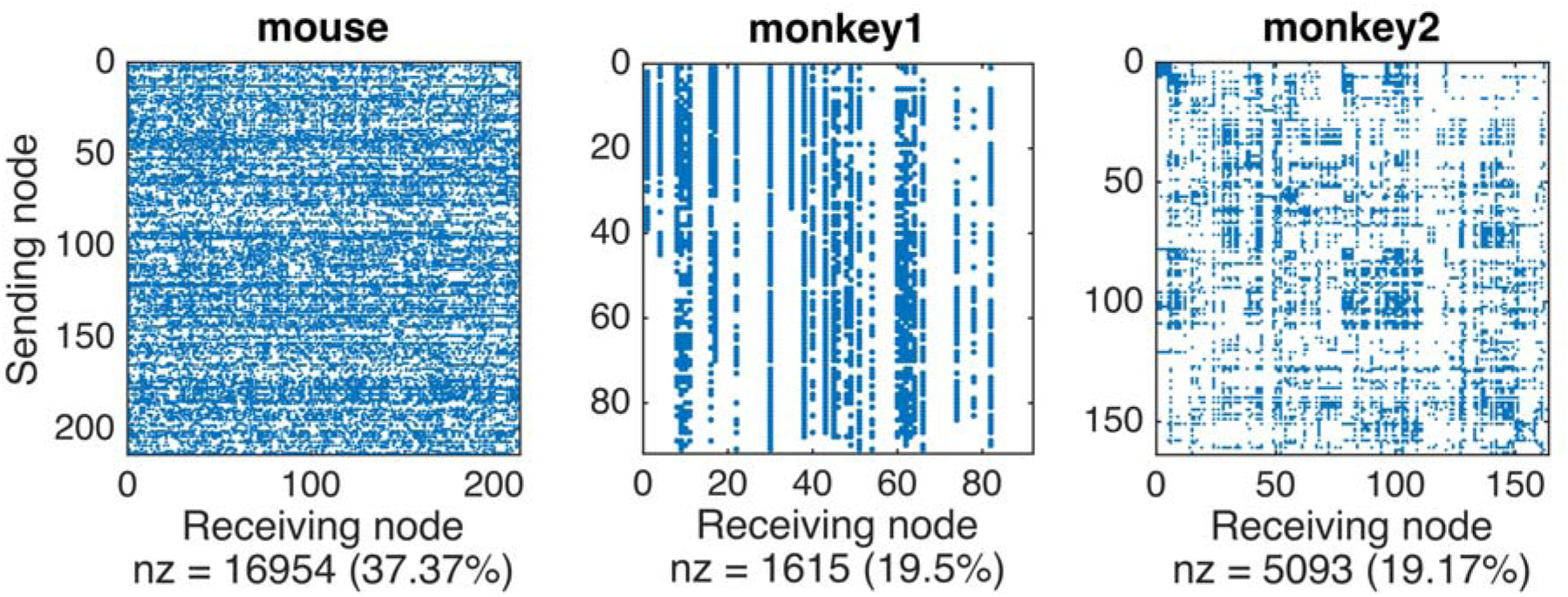
Adjacency matrices of the connectomes tested. *Mouse* includes the ipsilateral whole brain of the mouse (Oh et al., 2014), while *monkey1* (Markov et al., 2014) and *monkey2* (Bakker et al., 2012) include monkey cortex. All edges (blue dots) are directed and have weight 1. Non-zero entries are shown in numerical and percentage terms below each matrix.

### Message-passing Models

In our models, time is discretized and message-passing at each node is synchronous. During a time step, a node can be active or inactive (0 or 1). Each time step *t, L* new messages are injected into the system at randomly chosen nodes, representing load. Variable loads are to be expected in brain networks, but are most likely restricted to a relatively low range given metabolic constraints. Because the three connectomes have different numbers of nodes *N*, load is expressed as a percentage of the number of nodes in the network. We also report the case of 1 new message injected per time step in each case. We restrict simulations to the load regime with a number of messages less than or equal to 50% of *N*.

During each time step, a node looks to all incoming messages, including messages that are already in the system and new messages that are scheduled to be injected; if there is more than one message coming in, all are deleted. The full set of operations in the model is given as pseudocode in Supplemental Box 1. We are also making code to generate these models available on GitHub.

### Measures of Dynamic Network Activity

Simulations consist of 500 runs of 1000 time steps each. Data from the first 500 time steps is excluded to allow the analysis of equilibrium dynamics (see, e.g., Schrubin and Margolin, 1978). However, we note that data for the full test run produces the same overall pattern of results.

First, we measured the fraction of nodes active on a given time step. We measured both attempted activity before accounting for collisions, and actual (net) activity after collisions.

We also measured the sparseness over time in 5-time step windows (bins) using the Treves-Rolls measure of sparseness (Treves and Rolls, 1991; Rolls and Tovee, 1994); recall that sparseness here relates to network activity rather than to network structure. The Treves-Rolls measure is sensitive to higher-order statistical regularities in data (see Willmore and Tolhurst, 2001). We calculate population sparseness by constructing a histogram of the frequency of firing *x*_*n*_ within a time window per node, then sum over *i* nodes. The Treves-Rolls measure *S* (Equation 1) divides the square of the sum of the distribution by the sum of the squares of the distribution:

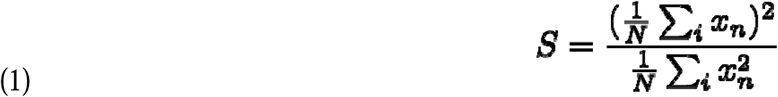

This measure ranges between maximal sparseness at *N*^*-1*^ if one unit is active per window, and all other units are inactive; minimal sparseness is 1.0 for uniformly-distributed activity (sparseness of standard normal distribution by this measure is 0.64). See Figure 2 for an illustration of this measure. Lower values of Treves-Rolls sparseness indicate more sparse activity. We note that tests of 10-time step windows for the sparseness measure produced similar results. We also calculated lifetime sparsene s as above, summing over time instead of over nodes.

**Figure 2.**
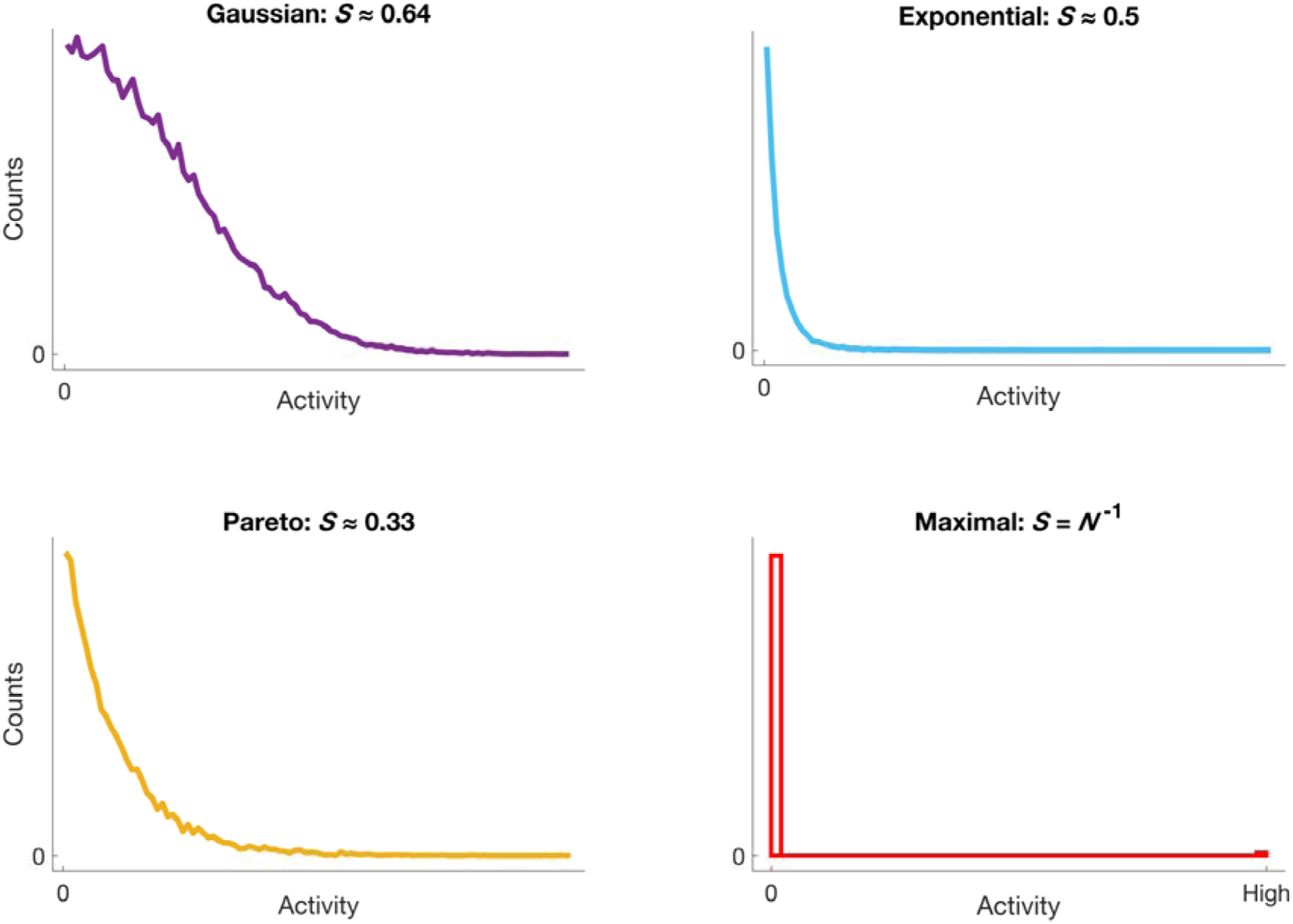
Illustration of the Treves-Rolls measure of sparseness *S* for hypothetical histograms of activity data. Shown are histograms that are well approximated by common functions, along with corresponding sparseness values as calculated from Equation 1. Top left: (half) Gaussian, *S* ≈ 0.64; Top right: Exponential, *S* ≈ 0.5; Bottom left: Generalized Pareto (shape parameter = 0.25), *S* ≈ 0.33; Bottom right: Maximal sparseness (closest to zero), *S* = *N*^*-1*^, is achieved for a histogram where *x*_*n*_ are all zero except for one non-zero entry, indicated here by “High.” A uniform histogram (not shown) has minimal sparseness of *S* = 1.0.

### Comparison to Randomized Networks

To investigate the dependency of network dynamics on the specific wiring patterns of the mammal connectome, we compared network activity under our models to the same models implemented on randomized networks. Randomized networks have the same distribution of in- and out-degree as the empirical networks. This amounts to shuffling the in- and outgoing edges of nodes in a given network (Maslov and Sneppen, 2002).

However, the nature of the connectomes in this study complicates the randomization process. When a network is dense (in a topological sense; except for the present discussion of randomization, all references to density and sparseness refer to network activity rather than topology), as in the *mouse* connectome and the 29-node core of *monkey1*, network topology changes little after randomization. We therefore thresholded the *mouse* connectome to make it less dense for randomization. Since edge weights in the *mouse* are known (corresponding to tracer volumes) we thresholded the network to exclude edges with a weight below 0.0136 (a value close to the modal weight, and one implied as meaningful by Oh et al., 2014), then randomized the resulting network. Thresholding of the *mouse* network reduced degree in a way that brought edge density to a level comparable to that of *monkey2* (but interestingly, the global topology changed relatively little after removing more than half of the edges, suggesting self-similarity). The density problem is compounded in *monkey1* because the full 91-node network is low-density as compared to the other two networks. Therefore, we have omitted randomization of *monkey1*; results of randomized network comparison in the monkey are for the *monkey2* connectome, which has intermediate connectivity. Note that each trial in the randomized network simulations corresponds to a different randomized network. Other randomization approaches, such as that of Tipnis et al. (2018), have used iterative search among randomized networks (based on the method of Maslov and Sneppen, 2002) to find those that differ most from the empirical network.

However, this approach has potential for bias since only a single randomized network is generated, whereas our method compares the empirical network to a large family of randomizations.

## Results

Qualitative differences in network dynamics are apparent in visualizations of node activity over time. In Figure 3, we show attempted and actual (net) activity through a representative simulation of the *mouse* connectome, with nodes shown along the vertical axis and the last 500 time steps of the simulation running from left to right (*L* = 10%).

**Figure 3.**
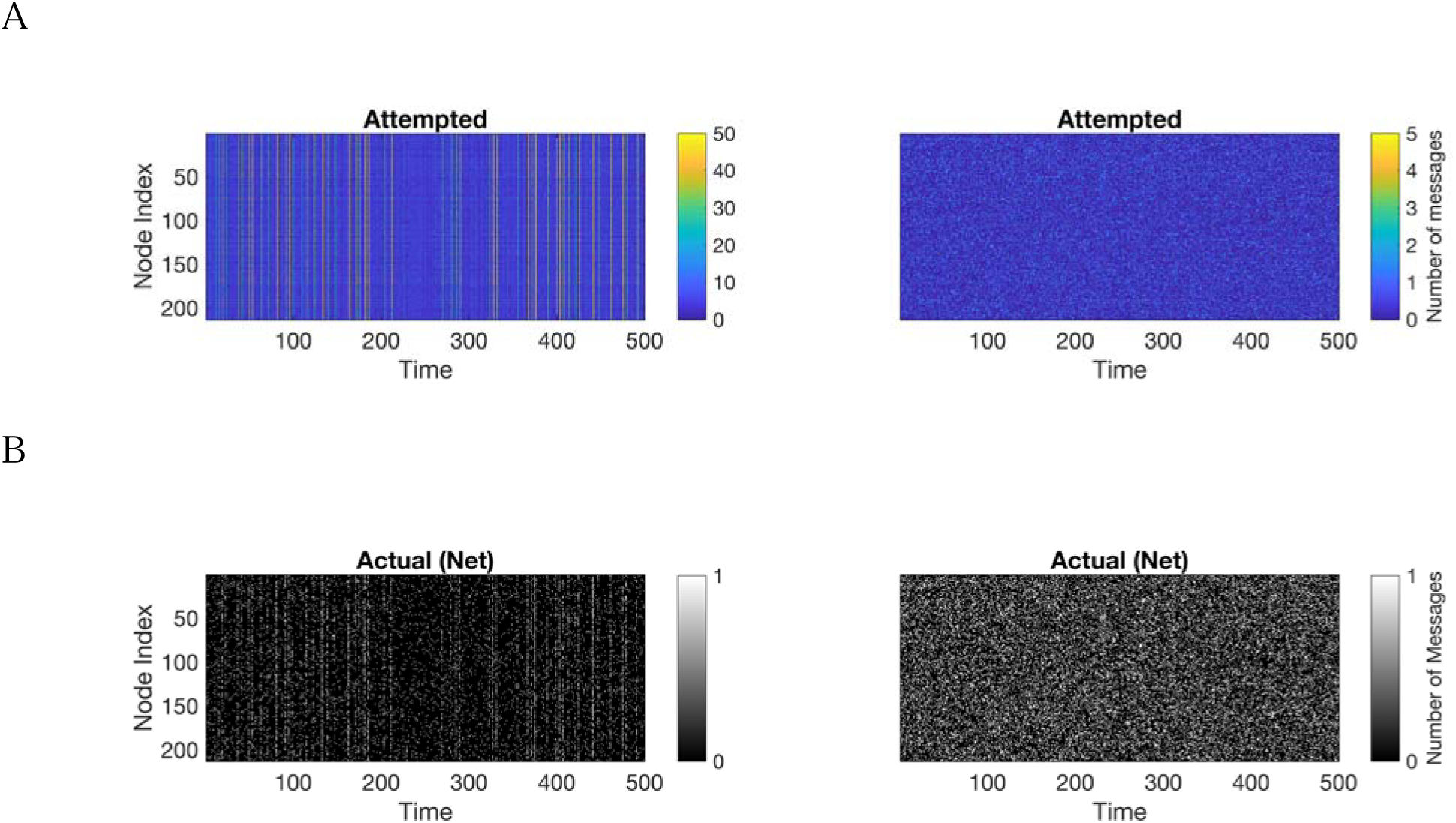
(A) Attempted activity (before collisions) and (B) actual activity (net, following collisions) under IS and RW models in the *mouse* connectome at 10% load. Vertical axis corresponds to nodes and time runs left to right. Colors indicated in the vertical axis in attempted activity plots (A) represent the number of colliding messages at a given node and time step. Net activity per node per time step (B) is binary (black=off, white=active).

Figure 3A shows attempted message-passing per node according to the color scale shown (i.e., the number of messages colliding). In general, attempted messages are distributed in unimodal, monotonically decreasing fashion in RW whereas attempted messages are bimodally distributed in IS with a high peak at zero (not shown).

Figure 3B shows actual activity for the IS model (left panel) and the RW model (right panel). From inspection, the net activity is more “bursty” and sparse in IS while activity is more uniformly distributed across nodes and time in RW.

We can quantify these observations by calculating the average net activity and the sparseness of activity across simulations. Figure 4 shows average net activity (expressed as a fraction of the number of nodes active in a given model, averaged over time and trials) as a function of load *L* (expressed as a percentage of nodes creating new messages on each time step, except for the left-most tick, which represents injection of one new message per time step). IS models are shown with solid lines and RW models are shown with dashed lines. Though RW models have lower net activity with *L* = 1 message, net activity increases rapidly and monotonically as load increases. The relationship is well-fit by a log function (not shown). In contrast, IS models show lower activity that is relatively uniform across load.

**Figure 4.**
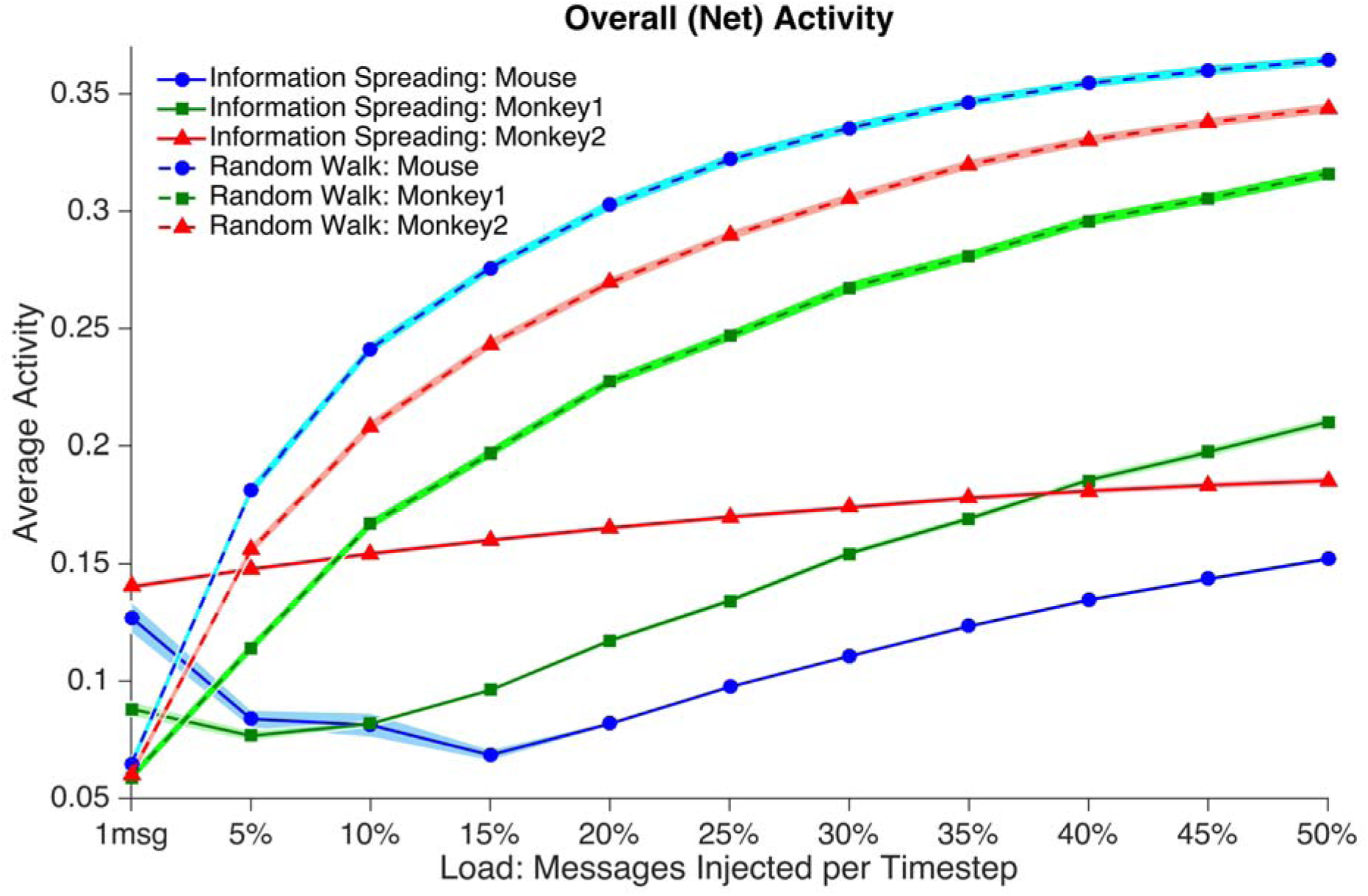
Mean of net simulated activity on the mammal connectome as a function of load under IS (solid lines) and RW (dashed lines) models for *mouse* (circles), *monkey1* (squares) and *monkey2* (triangles) networks. In IS, activity remains mostly constant across load and is lower compared to the corresponding value for RW at all load levels except 1 message per time step. Activity in RW increases by more than two octaves across loads tested. Shaded areas represent 2 standard deviations of variability from the mean across simulations (note that in some cases this dispersion is smaller than the width of the markers/lines). Data points show significant differences (t-test, p<0.01) for corresponding IS and RW measurements at all loads.

Figure 5 shows population sparseness of activity as a function of load. As with net activity across nodes, sparseness of activity in a given network is greater (closer to zero) for IS compared to RW at all loads except the minimum load. As a function of load, we observed the same pattern for sparseness over time within a simulation (lifetime sparseness of activity) as was seen for population sparseness, namely that sparseness over time in a given network is greater (closer to zero) for IS compared to RW except at the lowest loads. (See Supplemental Figure 1). We note that, like attempted activity, net activity over time for IS is bimodally distributed with a high peak at zero, whereas net activity under RW is distributed in roughly Gaussian fashion (See Supplemental Figure 2).

**Figure 5.**
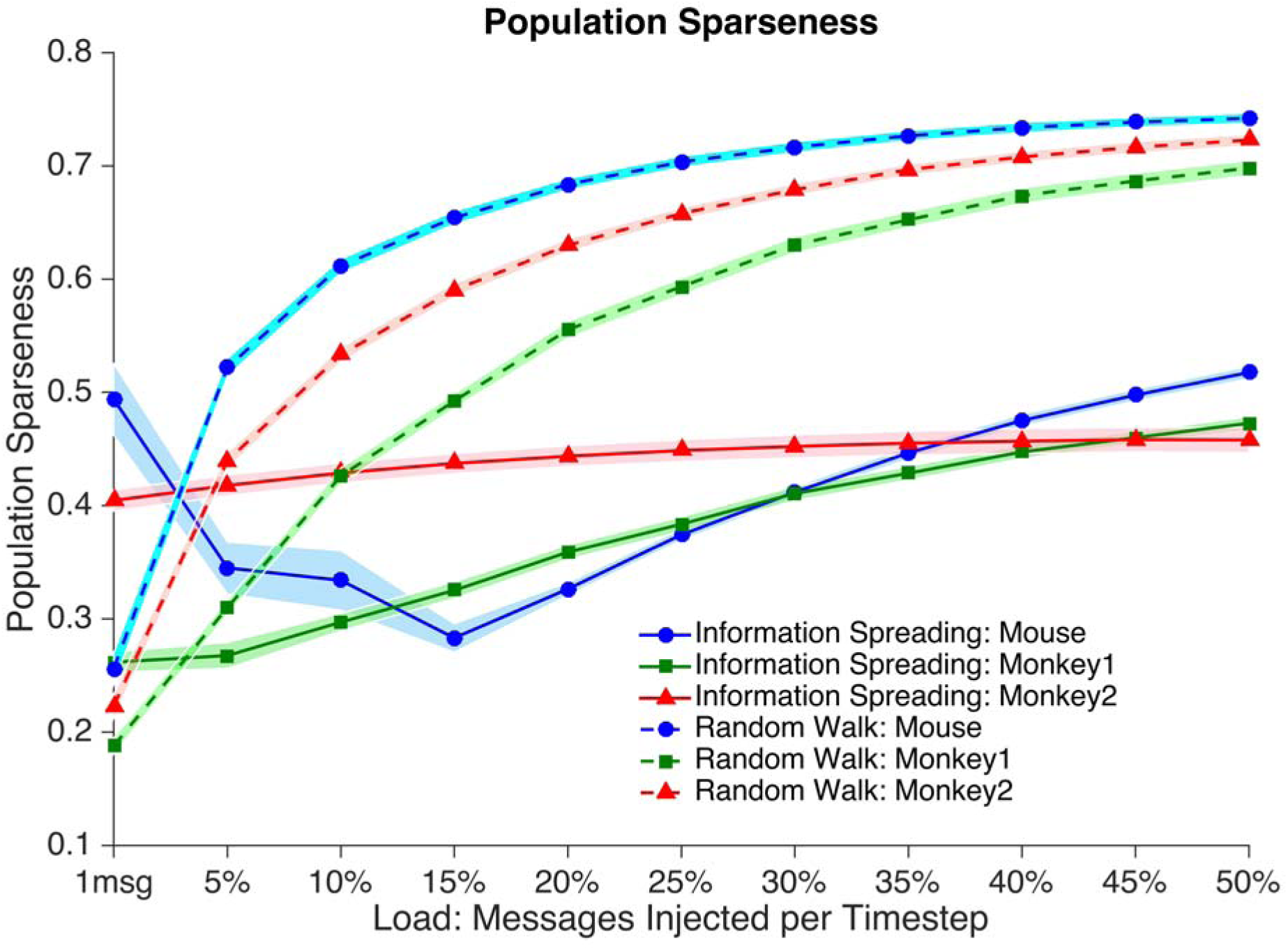
Population sparseness of activity as a function of load under IS (solid lines) and RW (dashed lines) models for *mouse* (circles), *monkey1* (squares) and *monkey2* (triangles) networks. As with net activity, IS models show relatively constant sparseness of activity across load, with greater sparseness of activity (closer to zero) than RW models at all loads except 1 message per time step. Sparseness in RW decreases by more than a factor of 2. Shaded areas represent 2 standard deviations of variability from the mean of population sparseness across simulations. Data points show significant differences (t-test, p<0.01) for corresponding IS and RW measurements at all loads.

Results of comparison to randomized networks (Figure 6) in *monkey2* and the thresholded *mouse* show that IS models have lower net activity and greater sparseness in the empirical network at almost all loads tested. RW models on empirical networks generally show very small decreases (*monkey2*) or slight increases in activity compared to randomized networks (thresholded *mouse*).

**Figure 6.**
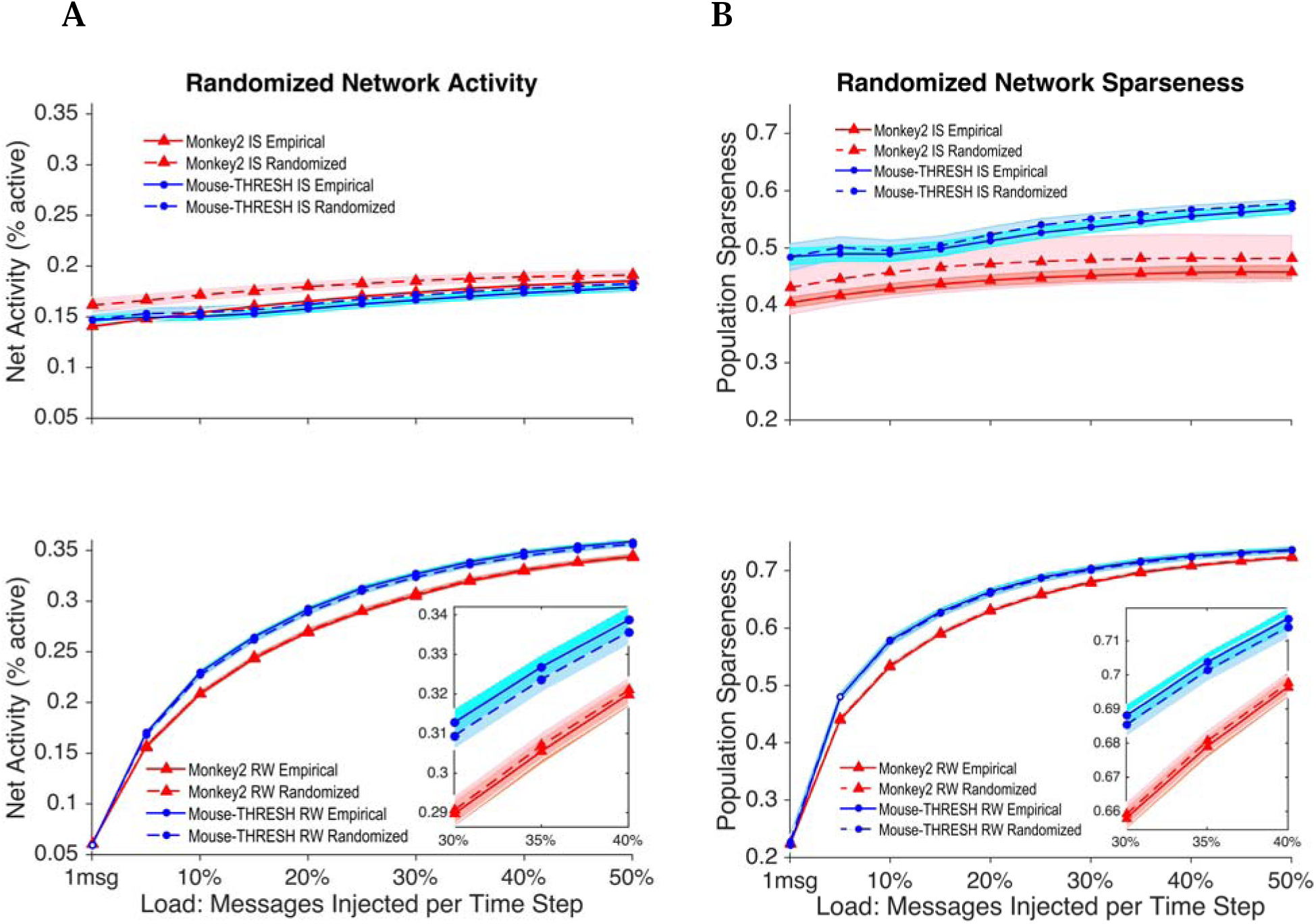
(A) Net activity and (B) population sparseness for empirical and randomized networks of the *monkey2* and the (thresholded) *mouse* under IS and RW models. In IS models, both the empirical *monkey2* network (triangles) and the (thresholded) *mouse* network (circles) show lower activity and greater sparseness (showing a difference in percentage terms of between 2-7%) compared to corresponding randomized networks. Randomized RW models show largely the same behavior as the empirical networks (differing by less than 1%) in terms of activity and sparseness. Filled symbols indicate significant differences (t-test, p<0.01) between empirical and corresponding randomized networks, whereas open symbols indicate no significant difference.

More specifically, empirical *monkey2* and thresholded *mouse* networks under IS differ significantly (t-test, p<0.01) at all loads compared to corresponding randomized networks. Under IS, the mean net activity of the empirical networks across loads is lower by 0.013 (*monkey2*) and 0.0039 (thresholded *mouse*) activity units compared to corresponding randomized networks. In percentage terms, this equates to a difference in activity between empirical and randomized networks of 7.3% and 2.3%, respectively. The mean sparseness of the empirical network across loads under IS is greater (closer to zero) by 0.026 (*monkey2*) and 0.0092 (thresholded *mouse*) units compared to corresponding randomized networks. This equates to a difference in sparseness of 5.6% and 1.7%, respectively. Empirical *monkey2* and thresholded *mouse* networks under RW also differ significantly (t-test, p<0.01) at all but 2 load values compared to corresponding randomized networks. However, compared to corresponding randomized networks, the mean net activity of the empirical networks across loads under RW is marginally lower by 0.00074 (*monkey2*) or greater by 0.0027 (thresholded *mouse*) activity units. This equates to a difference in net activity of 0.30% and 0.93%, respectively. Likewise, the mean sparseness of the empirical network across loads under RW is marginally greater by 0.0013 (*monkey2*) or lower (closer to one) by 0.0012 (*mouse*) units compared to corresponding randomized networks. This equates to a difference in sparseness of 0.25% and 0.027%, respectively.

We note that the same pattern of results also obtains on the unthresholded *mouse* network. Although the differences between empirical and randomized networks are not large, they are uniformly significant. This evidence is consistent with the notion that empirical network topology in mammals promotes sparse, efficient activity under an information spreading routing strategy.

## Discussion

In the context of collision interactions among signals, we used numerical simulations to show how the mammal brain can exploit mesoscale network structure by spreading information widely, and in particular by sending multiple copies of an incoming signal. Under an information spreading (IS) strategy for routing, the network achieves globally low net activity and high sparseness of activity like what is found in real mammal brains. Net activity and sparseness of activity change relatively little as load increases under the IS model. This result suggests a way that the brain could achieve ongoing sparse activity spread across the entire system using local protocol, but without deviating too far from globally homeostatic operation. In contrast, random walk (RW) models with the same collision rule produce substantially higher summed activity. In RW, substantial numbers of nodes are active in a given time window, resulting in less sparse activity. Net activity and sparseness of activity under RW change substantially as load increases over the same range, with activity increasing more than four-fold, and sparseness decreasing by around a factor of two. Our findings hold in both the monkey cortex and mouse whole brain. In addition, empirical networks under IS models show lower activity and greater sparseness of activity than randomized networks with the same degree distributions.

Together, these results suggest that nonlinear collision protocol and mammal connectome topology could promote sparse, efficient routing of communication traffic across the mammal brain network. This protocol is biologically plausible, and does not require centralized control.

However, one can ask: are RW and IS models really comparable to each other? It is certainly the case that the actual number of messages circulating on the network is different under RW and IS models. For example, a single message injected as load will—on the next time step, and in the absence of incoming messages—lead to only one outgoing message under RW, while in IS it will lead to *k* outgoing messages (where *k* is out-degree) under these circumstances.

Yet such nonlinear interactions are precisely our object of study. Rather than seeing RW models as a control relative to IS models, the two models should be seen as extreme ends of a spectrum of diffusion-like routing strategies with distributed protocol. This spectrum is orthogonal to the spectrum of pathfinding strategies that span from shortest paths to random walks (see Avena-Koenigsberger et al., 2019). At one end of the spectrum of routing strategies, we have the random walk approach that is employed extensively to model diffusion-like processes with a conservation law. At the other end of the spectrum are models that generate spreading dynamics (e.g., those used to model infectious disease).

Random walk models are in any case very common in network neuroscience so their collision dynamics are of inherent interest to the field. We have shown that RW models have clear limitations in terms of achieving sparse, low, and relatively uniform activity across loads given simple collision rules.

Taking a wider perspective, the management of signals arising from different sources that impinge on the same target is a critical problem in network neuroscience. In the mesoscale mammal connectome, where all nodes are only a few hops from each other (and where signals are excitatory), the system would seem to need a strategy for managing coincident excitations in order to route signals appropriately.

This conception stands in contrast to models of dynamics of the mesoscale connectome (particularly cortex) inspired by integrate-and-fire spiking neuron models, wherein the goal of the system is to have many excitatory signals arrive at the same time in order for the unit to be highly activated. Collisions— in the sense of multiple coincident efferents—are seen as signals to be integrated. However, studies such as Kaiser et al. (2007) and Kaiser and Hilgetag (2010) that approach dynamics from this perspective confirm the necessity of keeping global activity at a low and relatively constant level. Kaiser and colleagues tested a threshold routing model on surrogate hierarchical networks with biologically plausible parameters, and suggested that brain networks must seek regimes of “limited sustained activity.” Using a “spreading” model that is somewhat similar to our IS model, their simulations indicated that specific parameters of activity likelihood promoted limited sustained activity of around 10-20% of nodes. However, many if not most parameter settings led to “dying-out” activity or overload (Kaiser et al. 2007; Kaiser and Hilgetag, 2010). We have shown here that limited sustained activity of around 10-20% of nodes is robustly achieved on mammal connectomes under a nonlinear routing approach employing destructive collisions and information spreading. However, in contrast to Kaiser and Hilgetag (2010), our model shows that this pattern of activity is distributed across the entire network over time, rather than concentrated in a minority of nodes, and this behavior is demonstrated on all simulation runs.

In separate tests, we found that the structure of mammal brain networks is such that collisions that sum or threshold signals lead to very high activity and very low sparseness in both RW and IS models. This was suggested by tests of a “let one pass” collision rule whereby exactly one message in a collision of *m* messages is allowed to pass on a randomly chosen outgoing edge. Even at low load values, the network quickly overloads (especially under IS) with this collision rule producing data that are not meaningful, and therefore not reported here.

Our results show that strategies with nonlinear, self-organizing mechanisms that align with basic physiology can generate network activity that offsets collisions, requires no central agent, operates sparsely in time and over nodes, and maintains relatively uniform global activity despite large increases in signal load. However, automatic deletion is not the only or necessarily the best solution for managing collisions. There are many other possible strategies and combinations of strategies that could be at play including buffering (Mišić et al. 2014a), resending (Graham and Rockmore, 2011), and deflection routing (Fornito, Zalesky and Bullmore, 2016).

We find some evidence for one alternative strategy, which is to reduce load almost to zero. In this regime, redundancy may not be needed because collisions are rare, and a given path is unlikely to be congested. This is suggested in the finding that RW models achieve lower net activity and higher sparseness of activity than IS models at the lowest load (1 message per time step) in all cases. It remains possible that the brain could adopt this kind of extremely sparse strategy (see e.g., Ovsepian, 2015). However, such a system, being so close to floor values, would be restricted to low activity ranges and would be vulnerable to large changes in dynamics if load varies more than a small amount. In any case, if collisions are indeed very rare, this is also a major problem for models based on integrate-and-fire strategies and linear approaches.

We note that redundancy in the brain such as correlated firing among neighboring neurons (see e.g., Meytlis, Nichols and Nirenberg, 2012) is typically thought to be something that the system should minimize (Wainwright, 1999; Barlow, 2001). Following Shannon’s theory of information, parallel communication channels should seek to reduce mutual information to theoretical minima. However, the brain’s network topology is not parallel but rather small world-like (see, e.g., Achard et al., 2006; see also Knoblauch et al., 2016). In this situation, there is no straightforward way to apply Shannon’s theory for arbitrary node-to-node communication (El Gamal and Kim, 2011). Our results suggest that redundancy of signals may allow brain networks to strike a dynamic balance between signal creation and destruction.

However, copying all incoming signals has its drawbacks. More realistic solutions may be found between the extreme ends of the spectrum of routing strategies. For example, it may be more efficient to send redundant signals only on some fraction of outgoing edges. Tests exploring this parameter are underway; we predict a smooth transition in dynamics between the observed extremes of RW and IS conditions. In any case, the purpose of the present investigation is not to find the optimal solution for intrabrain communication but rather to address the question, how could a mammal brain efficiently route signals, given the likelihood of collisions, the need for distributed protocol, and a demand for low, sparse activity levels? Much as vision scientists have gained knowledge from asking what an efficient retinal code would be given the statistical regularities of the environment (e.g., Field, 1994; Graham and Field, 2009), we aim to advance model building by asking what an efficient whole-brain routing protocol would be given its architecture and operational demands.

We note that collisions cannot be ignored by assuming that signals arriving at a node at the same time do not interact. Of course, at the mesoscale, inputs arising in different nodes that travel to a target node do not necessarily synapse with the same population within the target node. But if there were dedicated links from sending nodes via a target node to destinations one synapse away, this would imply a different network topology (i.e. the target node would not be considered a node). If the brain performs dynamic routing at nodes—and functional demands appear to necessitate this (Olshausen et al., 1993; Graham and Rockmore, 2011)—collisions must be managed.

Nevertheless, the brain probably does not annihilate all colliding signals at all nodes. The rules governing collisions may vary in time and space from those that are fully punitive for coincident signals to those that require greater coincidence. This is a limitation of our study. Collision rules are a dimension worth exploring but as an initial study, we adopted a simplified model that promotes interpretability.

The difficulty of studying nonlinear interactions of brain network communication is due in part to the problem of simulating dynamics explicitly. The challenge lies in tracking every signal’s path and interactions. Random walk models and shortest path models admit analytical solutions that capture the likelihood of a set of states of the network. But this is generally possible only when collisions and other nonlinear interactions are ignored. As a result, explicit models capable of accounting for collisions have been relatively little studied to date (see Graham and Hao, 2018). With further study of explicit models we may be able to understand fine-grained activity patterns such as regional differences and correlations in activity. However, though we were able to model signal interactions explicitly, the lack of tracking of messages is a limitation of the present study. For example, we would like to know how many messages are delivered, and how quickly. We have identified a way to infer message “age” in our simulations and will present results of this work in a subsequent report.

This study has other limitations. One is the idiosyncrasies of the connectomes studied. For example, the CoCoMac network is known to be incomplete (Chen et al., 2020), and the Markov et al. (2014) data are limited by retrograde tracing procedures. In addition, we have not constrained or tailored activity in terms of functional demands.

Finally, we are limited in terms of the interpretation of our network randomization results. As noted earlier, randomizing networks in which nodes have a degree that is a significant fraction of the number of nodes leads to random networks that share edges with their empirical counterparts. However, in preliminary studies of the *monkey2* network, we found that a non-degree-preserving random network (an Erdős–Rényi network with the same number of nodes and edges, but where no edges are shared between empirical and randomized networks) shows significantly higher activity and lower sparseness compared to empirical networks under both IS and RW. In fact, the difference between empirical and randomized networks is somewhat more pronounced compared to the degree-preserving case, indicating that degree distributions do play a role in generating low, sparse activity, but they do not fully explain this effect.

We conclude that the mammal brain network appears capable of striking a balance between activity and collisions such that sparse activity is robustly maintained across a wide range of loads. At present we cannot say what properties of network structure are necessary for supporting this behavior. Indeed, it remains unknown whether the “creative destruction” we observe here is unique to brains or constitutes a more widespread behavior of complex networks. Nevertheless, we argue that some strategy for managing collisions is necessary, and whatever solution the mammal brain has adopted, it must generate low, stable net activity that is sparsely distributed across the network.

## Acknowledgments

We thank Barbara Finlay, David Field, Marc-Thorsten Hütt, and Ao Tang for helpful discussions.

## Author Contributions

YH and DG conceptualized the study, designed the framework, performed analyses, interpreted the results, and wrote the manuscript.

## Code Availability

Matlab code for performing simulations under IS and RW models will be posted to GitHub.

## Supplemental Material

**Box S1.**
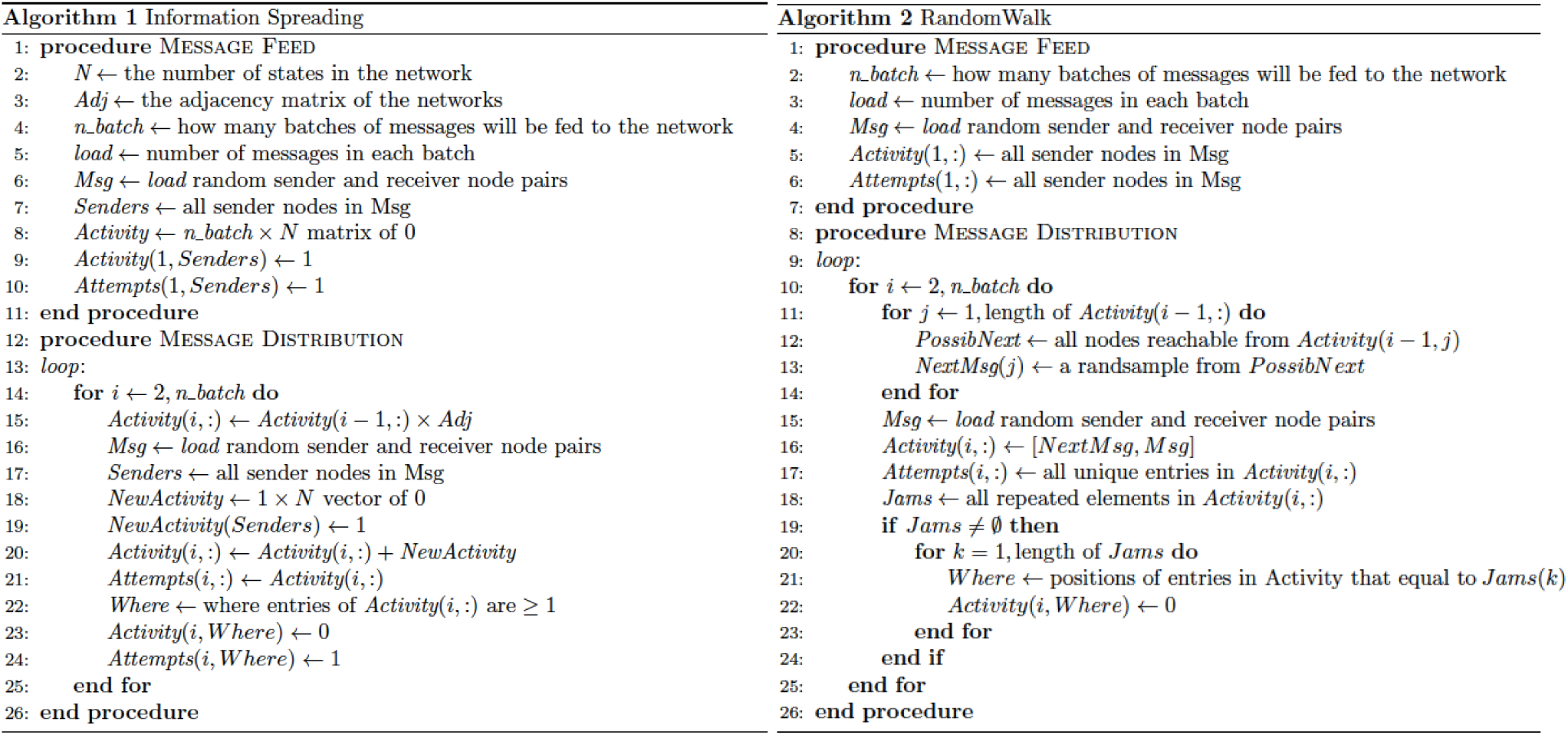
(A) Pseudocode of information spreading (IS) model. (B) Pseudocode of random walk (RW) model.

**Figure S1.**
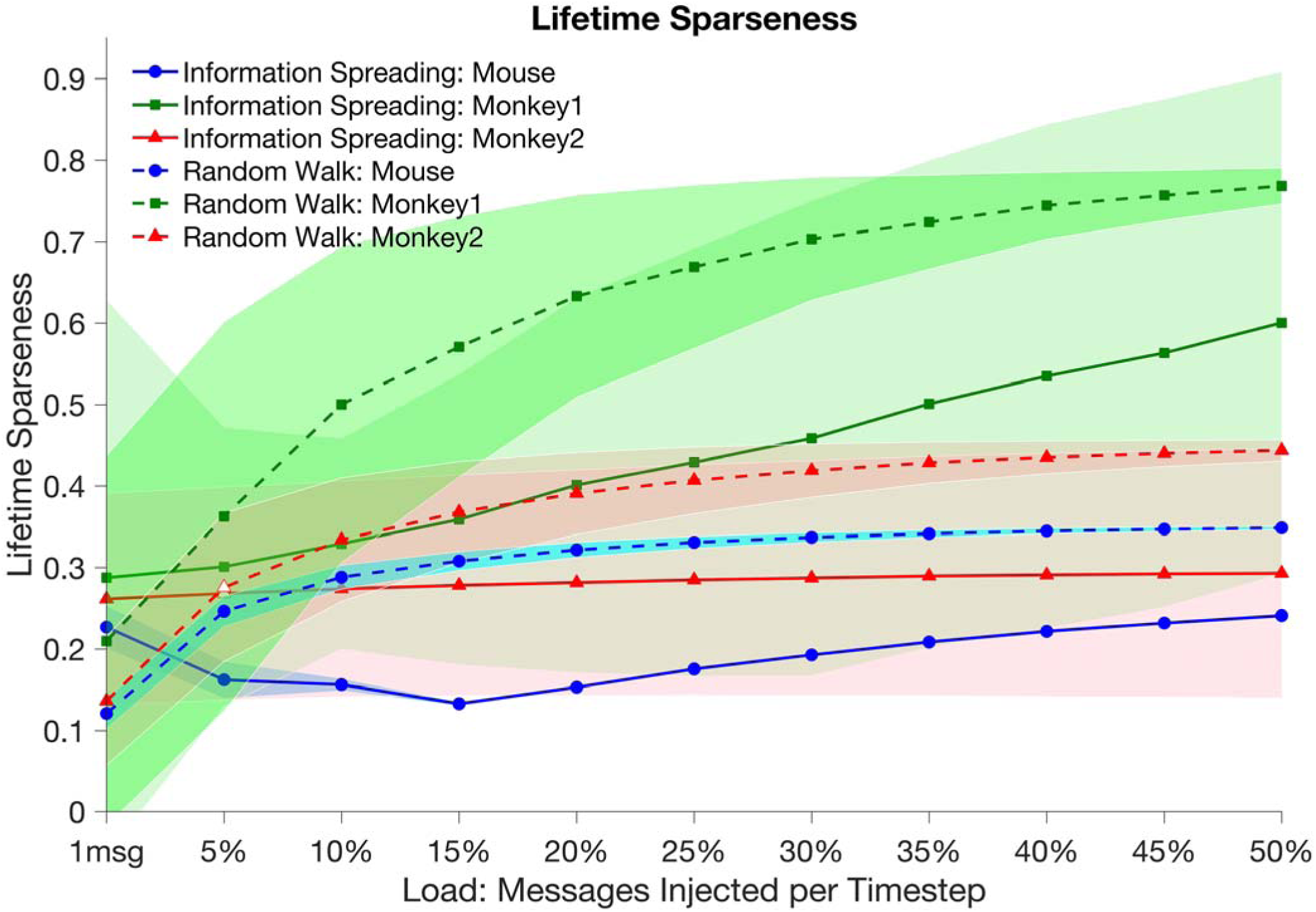
Mean lifetime sparseness of IS (solid lines) and RW (dashed lines) models as a function of load. As with population sparseness, lifetime sparseness is greater (closer to 0) for IS models compared to RW at loads above 1 message per time step. Shaded areas indicate 2 standard deviations from the mean. Filled symbols indicate significant differences (t-test, p<0.01) between corresponding IS and RW data, whereas open symbol indicates no significant difference.

**Figure S2.**
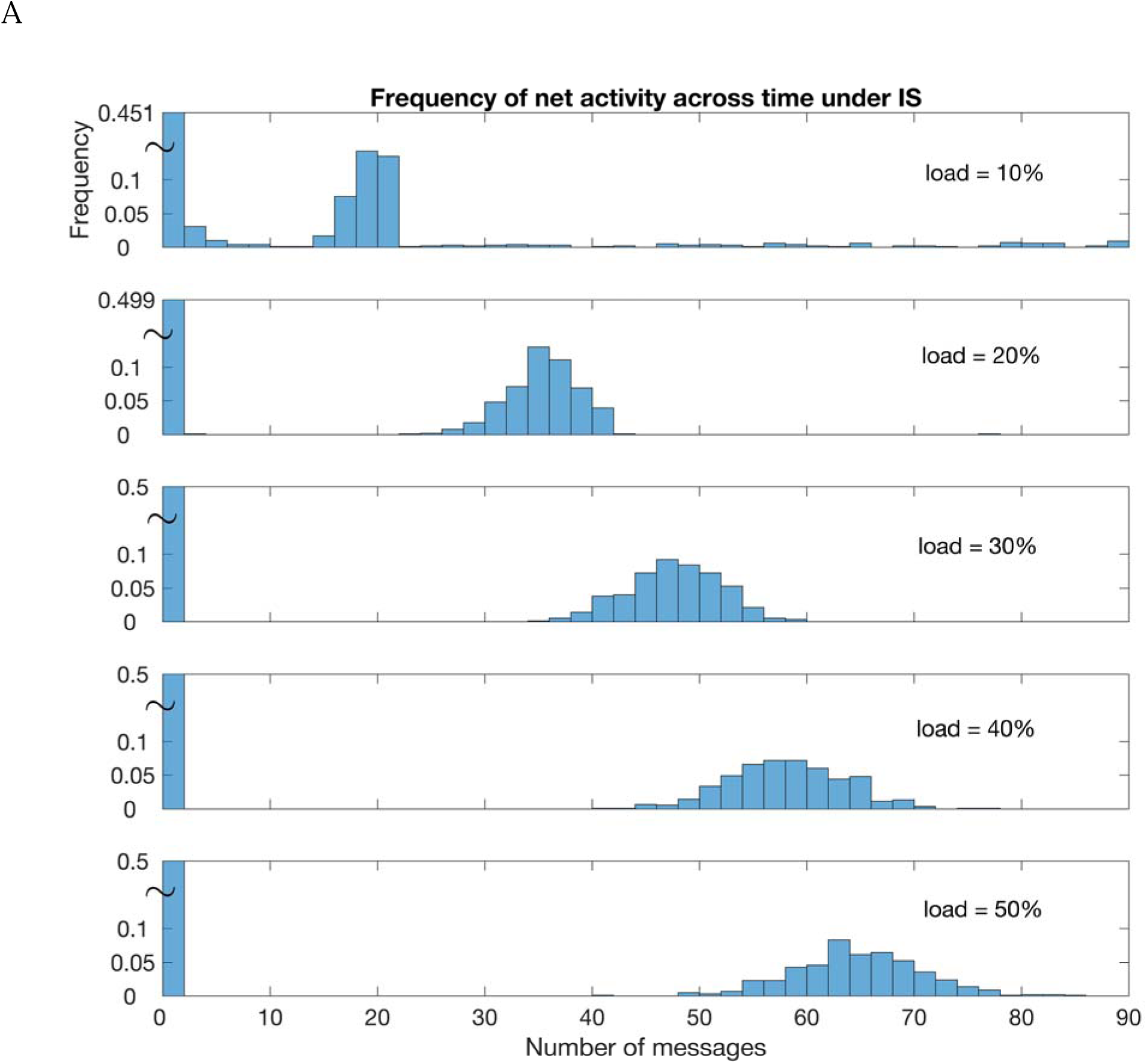
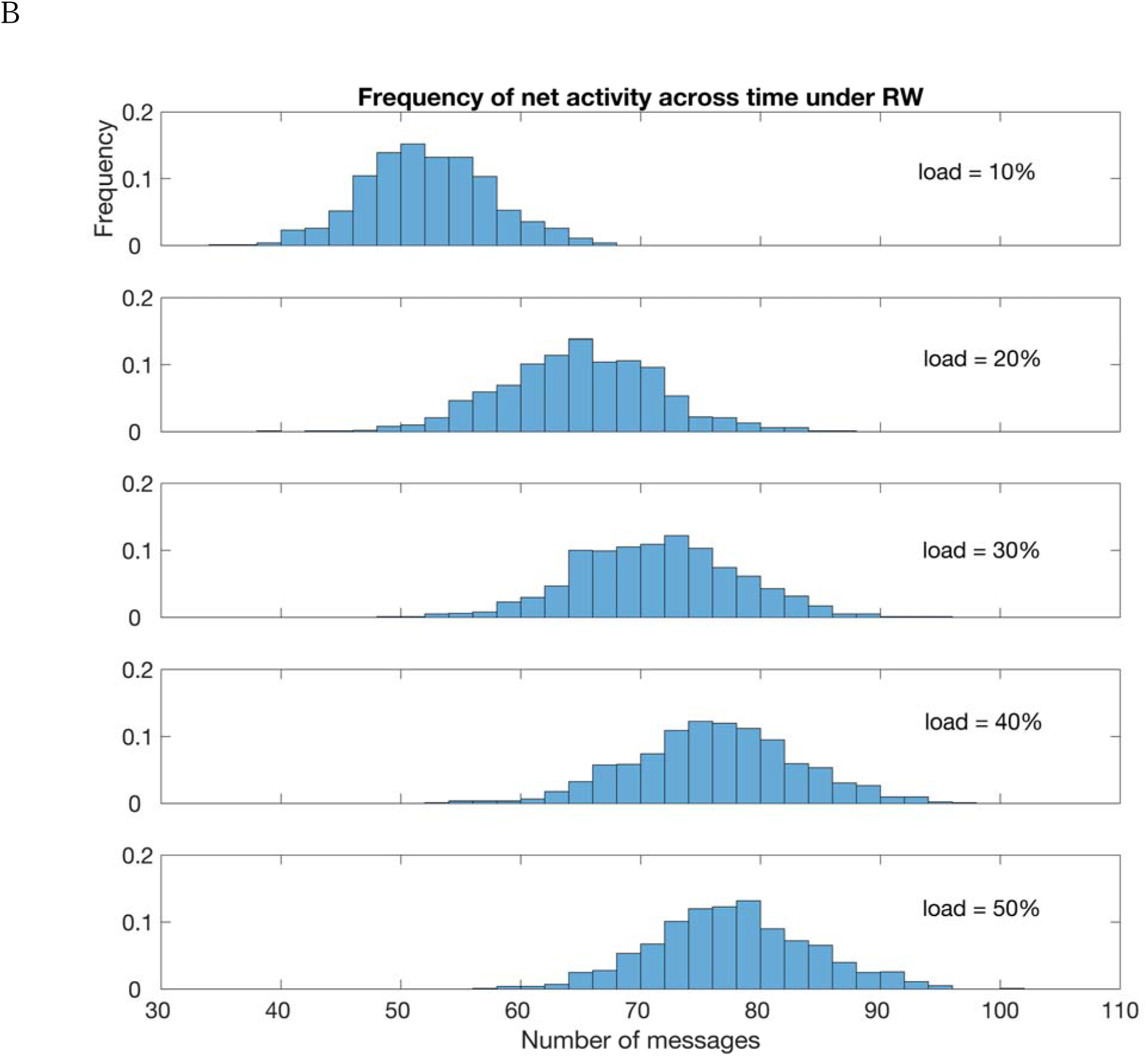
Relative frequency of net activity under IS models (A) and under RW models (B) at different loads. Results are for the *mouse* network. Tildes in A indicate a contracted vertical axis.

